# Discovery of biomarkers for glycaemic deterioration before and after the onset of type 2 diabetes: an overview of the data from the epidemiological studies within the IMI DIRECT Consortium

**DOI:** 10.1101/300244

**Authors:** Robert W. Koivula, Ian M. Forgie, Azra Kurbasic, Ana Viñuela, Alison Heggie, Giuseppe N. Giordano, Tue H. Hansen, Michelle Hudson, Anitra Koopman, Femke Rutters, Maritta Siloaho, Kristine H. Allin, Søren Brage, Caroline A. Brorsson, Adem Y. Dawed, Federico De Masi, Christopher J. Groves, Tarja Kokkola, Anubha Mahajan, Mandy H. Perry, Simone P. Rauh, Martin Ridderstråle, Harriet J. A. Teare, Louise Thomas, Andrea Tura, Henrik Vestergaard, Tom White, Jerzy Adamski, Jimmy Bell, Søren Brunak, Jacqueline Dekker, Emmanouil T. Dermitzakis, Philippe Froguel, Gary Frost, Ramneek Gupta, Torben Hansen, Andrew Hattersley, Bernd Jablonka, Markku Laakso, Timothy J. McDonald, Oluf Pedersen, Jochen M. Schwenk, Imre Pavo, Andrea Mari, Mark I. McCarthy, Hartmut Ruetten, Mark Walker, Ewan Pearson, Paul W. Franks, for the IMIDIRECT Consortium

## Abstract

**Background and aims:** Understanding the aetiology, clinical presentation and prognosis of type 2 diabetes (T2D) and optimizing its treatment might be facilitated by biomarkers that help predict a person’s susceptibility to the risk factors that cause diabetes or its complications, or response to treatment. The IMI DIRECT (Diabetes Research on Patient Stratification) Study is a European Union (EU) Innovative Medicines Initiative (IMI) project that seeks to test these hypotheses in two recently established epidemiological cohorts. Here, we describe the characteristics of these cohorts at baseline and at the first main follow-up examination (18-months).

**Materials and methods:** From a sampling-frame of 24,682 European-ancestry adults in whom detailed health information was available, participants at varying risk of glycaemic deterioration were identified using a risk prediction algorithm and enrolled into a prospective cohort study (n=2127) undertaken at four study centres across Europe (Cohort 1: prediabetes). We also recruited people from clinical registries with recently diagnosed T2D (n=789) into a second cohort study (Cohort 2: diabetes). The two cohorts were studied in parallel with matched protocols. Endogenous insulin secretion and insulin sensitivity were modelled from frequently sampled 75g oral glucose tolerance (OGTT) in Cohort 1 and with mixed-meal tolerance tests (MMTT) in Cohort 2. Additional metabolic biochemistry was determined using blood samples taken when fasted and during the tolerance tests. Body composition was assessed using MRI and lifestyle measures through self-report and objective methods.

**Results:** Using ADA-2011 glycaemic categories, 33% (n=693) of Cohort 1 (prediabetes) had normal glucose regulation (NGR), and 67% (n=1419) had impaired glucose regulation (IGR). 76% of the cohort was male, age=62(6.2) years; BMI=27.9(4.0) kg/m2; fasting glucose=5.7(0.6) mmol/l; 2-hr glucose=5.9(1.6) mmol/l [mean(SD)]. At follow-up, 18.6(1.4) months after baseline, fasting glucose=5.8(0.6) mmol/l; 2-hr OGTT glucose=6.1(1.7) mmol/l [mean(SD)]. In Cohort 2 (diabetes): 65% (n=508) were lifestyle treated (LS) and 35% (n=271) were lifestyle + metformin treated (LS+MET). 58% of the cohort was male, age=62(8.1) years; BMI=30.5(5.0) kg/m2; fasting glucose=7.2(1.4)mmol/l; 2-hr glucose=8.6(2.8) mmol/l [mean(SD)]. At follow-up, 18.2(0.6) months after baseline, fasting glucose=7.8(1.8) mmol/l; 2-hr MMTT glucose=9.5(3.3) mmol/l [mean(SD)].

**Conclusion:** The epidemiological IMI DIRECT cohorts are the most intensely characterised prospective studies of glycaemic deterioration to date. Data from these cohorts help illustrate the heterogeneous characteristics of people at risk of or with T2D, highlighting the rationale for biomarker stratification of the disease - the primary objective of the IMI DIRECT consortium.

**Abbreviations:**

ASAT: Abdominal subcutaneous adipose tissue
DIRECT: Diabetes Research on Patient Stratification
EU: European Union
MMTT: Mixed-meal tolerance test
MRI: Magnetic resonance imaging
hpfVM: High-pass filtered vector magnitude
IAAT: Intra-abdominal adipose tissue
IGR: Impaired glucose regulation
IMI: Innovative Medicines Initiative
ME: multiecho
NGR: Normal glucose regulation
OGTT: Oral glucose tolerance test
PA: Physical activity
TAAT: Total abdominal adipose tissue
T2D: Type 2 Diabetes

## Introduction

The global prevalence of type 2 diabetes (T2D) is burgeoning. There is no cure, nor are there treatments effective enough to halt the progression of the disease. The burden the disease conveys at a societal and personal level are enormous, with an estimated prevalence of around 415 million people (8.8% of adults) globally in 2015 [1]. In Europe alone, the cost of diagnosing and treating the disease and its complications in 2015 was estimated to be around €140 billion [1]. This bleak picture emphasises the profound shortcomings in our understanding of T2D aetiology and pathogenesis, and the inadequate tools available with which to combat the disease.

Like some other complex diseases, the clinical presentation and prognosis of T2D is heterogeneous. The risk conveyed by established diabetogenic factors such as obesity, physical inactivity and certain dietary components varies widely from one person to the next, as does their response to interventions targeting these risk factors. This is also true for those in whom diabetes is manifest, with response to anti-diabetic therapies, occurrence of adverse events and rates of progression being variable and hard to predict.

The diagnosis of T2D is relatively straightforward, relying primarily on evidence of chronically elevated blood glucose concentrations [2]. However, elevated blood glucose concentrations can be the consequence of multiple defects in energy metabolism occurring across several organs and tissues [3–5] caused by myriad acquired or inherited factors. Thus, T2D as currently defined, characterises a collection of underlying pathologies [6], each with the common feature of elevated blood glucose, that may require tailored therapies. The stratification of T2D into treatable subclasses might be possible if accessible biomarkers of the disease’s underlying pathologies were known.

Although improving the management of T2D through sub-classification may lead to more focused treatment, susceptibility to risk factors and response to treatments also vary. Thus, stratifying patient populations into subgroups defined using biomarkers quantifying susceptibility to risk factors and responsiveness to specific therapeutics would further enhance our ability to treat, and ideally prevent, the disease.

The IMI DIRECT consortium is a collaboration among investigators from some of Europe’s leading academic institutions and pharmaceutical companies [7]. The overarching objective of IMI DIRECT is to discover and validate biomarkers of glycaemic deterioration before and after the onset of T2D. To this end, we established two new multicentre prospective cohort studies comprised of adults from northern Europe at risk of, or with, recently diagnosed T2D. Within these cohorts, a comprehensive array of risk factors, intermediate phenotypes, and metabolic outcomes are repeatedly assessed using cutting-edge technologies.

Here we: i) describe the characteristics of the two IMI DIRECT cohorts at baseline and at the first major follow-up visit 18-months later, to provide context for those subsequently analysing and reviewing these data; and ii) consider these results in the context of the implemented protocols and plans outlined at the beginning of the project [7].

## Methods

The rationale and design of the epidemiological cohorts within IMI DIRECT are reported elsewhere [7]; here we provide data and information about key variables and methods respectively that are not previously described.

Approval for the study protocol was obtained from each of the regional research ethics review boards separately and all participants provided written informed consent at enrolment. The research conformed to the ethical principles for medical research involving human subjects outlined in the declaration of Helsinki.

### Recruitment, enrolment and eligibility

The derivations of Cohort 1 and Cohort 2 are shown in Figure 1. The sampling-frame for Cohort 1 was derived from four existing prospective cohort studies: *Metabolic Syndrome in Men* (METSIM) study (Finland) [8]; *Relationship between Insulin Sensitivity and Cardiovascular disease* (RISC) [9], *Hoorn Meal Study* (HMS), and *New Hoorn Study* (NHS) [10] (Netherlands); *Health2010* [11], *Health2006* [12], *Danish Study of Functional Disorders* (DanFunD) [13], and *Gut*, *Grain and Greens* (GGG) [14] studies (Denmark); *Malmö Diet and Cancer* (MDC) study (Sweden) [15]. Participants for Cohort 2 were identified through general practice and other registries, as described previously [7].

**Figure 1.**
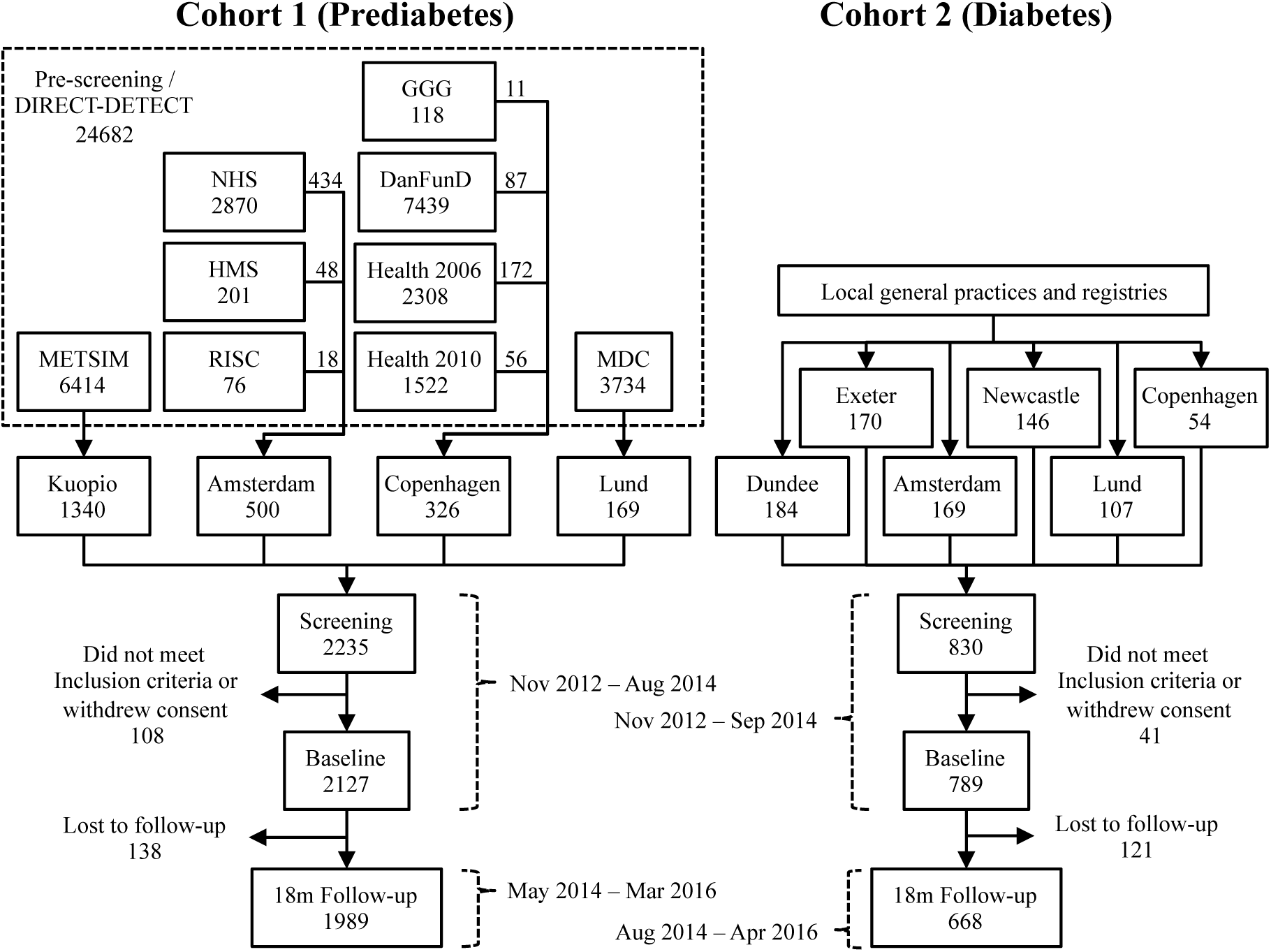
Participant flow of Cohorts 1 and 2. METSIM: Metabolic Syndrome in Men study, NHS: New Hoorn Study, HMS: Hoorn Meal Study, RISC: Relationship between Insulin Sensitivity and Cardiovascular disease cohort, GGG: Gut, Grain and Greens study, DanFunD: Danish Functional Disability study.

After excluding participants who did not meet the inclusion criteria or whose data failed quality control, a total of 2127 and 789 participants at risk of or with T2D were retained in Cohort 1 and Cohort 2, respectively. For Cohort 1, emphasis was placed on recruiting participants deemed at high-risk of T2D, i.e. those with HbA1c concentrations between 37-48 mmol/mol (5.5-6.5%). As anticipated during the design phase, the sampling-frame contained too few participants that fulfilled this criteria; thus, we proceeded to enrol participants with progressively lower HbA1c concentrations, who were also considered at highest risk of glycaemic deterioration, based on the DIRECT-DETECT risk algorithm applied to the full sampling-frame [7, 16].

In Cohort 1, 1989 (93%) participants enrolled at baseline also attended their first major follow-up visit at 18.6(1.4) months [mean(SD)]. Similarly in Cohort 2, 668 (85%) participants enrolled at baseline attended their first major follow-up visit at 18.2(0.6) months [mean(SD)].

### Glycaemic biochemistry assays

Plasma glucose, insulin and C-peptide assays for Cohort 1 were carried out centrally at the University of Eastern Finland (Kuopio, Finland), where plasma glucose was analyzed using the enzymatic glucose hexokinase method and photometric measurement on Konelab 20 XT Clinical Chemistry analyzer (Thermo Fisher Scientific, Vantaa, Finland). In Cohort 2, plasma insulin and C-peptide were analysed using chemiluminometric immunoassay, (CLIA), Liaison Insulin and Liaison C-peptide (DiaSorin S.p.A, Saluggia, Italy). The instrument used was DiaSorin Liaison Analyzer, DiaSorin Deutschland GmbH, Dietzenbach, Germany. Plasma glucose, insulin and c-peptide assessments for Cohort 2 were carried out centrally at the University of Exeter (Exeter, UK). Assessments of Hba1c, blood lipids, ALT and AST for both cohorts were carried out centrally at the University of Exeter. Glucose was measured by enzymatic colorimetric assay GOD-PAP. Insulin was measured by electrochemiluminescence. C-peptide in plasma and urine was measured by electrochemiluminescence. HbA1c was meaured by ion-exchange High Performance Liquid Chromatography.

### Blood lipid and liver enzyme biochemistry assays

Triglyceride was measured by quantitative determination with glycerol blanking. HDL cholesterol was measured directly using PEG-modified enzymes and dextran sulfate. When cholesterol esterase and cholesterol oxidase enzymes are modified by PEG, they show selective catalytic activities toward lipoprotein fractions, with the reactivity increasing in the order: LDL < VLDL ≈ chylomicrons < HDL. Total cholesterol was measured by an enzymatic, colorimetric method. LDL cholesterol was calculated from the total cholesterol, HDL cholesterol and triglyceride concentrations using the Friedewald Formula: LDL = total cholesterol - HDL cholesterol - (triglyceride/2.2). ALT and AST were measured by UV absorbance without pyridoxal phosphate activation. ALT, AST, Cholesterol, Glucose, Triglycerides, and HDL cholesterol were measured using a Roche MODULAR P analyser (Roche Diagnostics, Inc, IN, USA). Insulin and C-peptide were measured using a Roche E170 analyser (Roche Diagnostics, Inc, IN, USA). HbA1c was measured using a Tosoh G8 analyser (Tosoh Bioscience, Inc, CA, USA).

### Blood GLP-1 assays

Plasma concentrations of GLP-1 levels were determined by drawing blood samples at two different time points (0 and 60 min) at the baseline visit during the 75g fsOGTT/MMTT. P800 tubes (Becton Dickenson, UK) were used to provide immediate protection from intrinsic proteolysis. Quantitative determination of active GLP-1 was achieved using MSD GLP-1 active kit (Meso Scale Diagnostics LLC, MD, USA. product code = K150JWC). Total GLP-1 was assayed using MSD GLP-1 total kit (Meso Scale Diagnostics LLC, MD, USA. product code = K150JVC).

### Abdominal MRI analyses

The volume of adipose tissue was measured in litres using magnetic resonance imaging (MRI), as described elsewhere [17]. *Total abdominal adipose tissue* (TAAT) may be separated into *intra*-*abdominal adipose tissue* (IAAT – also known as ‘visceral’ fat) and *abdominal subcutaneous adipose tissue* (ASAT). IAAT is the volume in litres of adipose tissue within the abdominal cavity. TAAT is the sum of IAAT and ASAT in litres. Liver and pancreas fat and iron (T2^∗^) were derived simultaneously using a multiecho (ME) MRI technique, as previously described [17, 18]. This method has the advantage over single voxel MRS in that regional differences in ectopic fat distribution can be measured. Furthermore, it is often possible to obtain a single slice quantification of the liver and pancreas, allowing simultaneous measurement of fat and iron within two separate organs. A biexponential curve-fitting model was used to derive the relative signal contributions from fat and water from the many images normally obtained with the ME sequence. Briefly, tissue with no fat infiltration generates a very smooth decay curve, whereas tissue containing a higher level of fat report significant oscillations throughout the decay curve [18]. A further output from the ME technique is T2^∗^ tissue values; as changes in these are indicative of iron content, it provides a clinically relevant additional measurement. Tissue iron concentration (Fe, units mg/g dry weight tissue) was estimated from T2^∗^ using a validated model [19].

### Diet assessment

Dietary intake was assessed by a 24-hour multi-pass method, using food habit and 24-hour recall questionnaires. Analysis of diet data was undertaken using Dietplan-6 (version 6.70.43, 2013; Forestfield Software, Horsham, UK). The analysis methods for these data are described in detail elsewhere [7].

### Physical activity assessment

Objective measures of physical activity (PA) were derived from triaxial accelerometers (ActiGraph GT3X+/GT3X+w/GT3X+bt, ActiGraph Co, Pensacola, USA) as described previously [7]. Raw data files (.gt3x) were converted to comma separated value (.csv) format storing rawest possible accelerations for each axis at a resolution of 30Hz using ActiLife 6 (version 6.11.5, ActiGraph Co, Pensacola, USA). All inferred measures of PA were calculated using PAMPRO (version uploaded 2015-10-21, MRC Epidemiology unit, Cambridge, UK), a custom open source software available under public license (https://github.com/Thomite/pampro). Data from each axis of acceleration was auto-calibrated to local gravity. Non-wear was inferred as a vector magnitude standard deviation of less than 4mGs for a consecutive period greater than 60-min. All measures presented here have been adjusted for diurnal rhythm to account for bias from non-wear removal. However, due to the wear method (non-dominant wrist fastened using the manufacturer non-removable hospital band), intermittent non-wear time was rare. The main PA estimates presented here are high-pass-filtered vector magnitude (hpfVM), which infers intensity of participants’ movement in any direction at any given time (here, averaged during wear period). Time spent in established physical activity intensities by physical activity energy expenditure was estimated using calculated hpfVM cutpoints (Sedentary: (<48 mGs hpfVM, Light: 48-154 mGs hpfVM, Moderate: 154-389 mGs hpfVM, Vigorous: >389 mGs hpfVM). The methods used to infer these measures have been validated and described in detail elsewhere [20].

### DNA Extraction and Genotyping

DNA extraction was carried out using Maxwell 16 Blood DNA purification kits and a Maxwell 16 semi-automated nucleic acid purification system (Promega). Genotyping was conducted using the Illumina HumanCore array (HCE24 v1.0) and genotypes were called using Illumina’s GenCall algorithm. Samples were excluded for any of the following reasons: call rate <97%; low or excess mean heterozygosity; gender discordance; and monozygosity. Genotyping quality control was then performed to provide high-quality genotype data for downstream analyses using the following criteria: call rate <99%; deviation from Hardy-Weinberg equilibrium (exact p<0.001); variants not mapped to human genome build GRCh37; and variants with duplicate chromosome positions (a total of 30,318 markers were excluded). A total of 3,032 samples and 517,958 markers across the two studies passed quality control procedures. We took autosomal variants with MAF>1% that passed quality control and constructed axes of genetic variation using principal components analysis implemented in the GCTA software to identify ethnic outliers defined as non-European ancestry using the 1000 Genomes samples as reference. We identified six individuals as ethnic outliers.

### Additional measures (not presented here)

Additional measures that are not described here include transcriptomics (RNA sequencing from fasting whole blood), microbiomics (DNA isolation and deep sequencing in faecal samples), proteomics (targeted array in fasting plasma) and metabolomics (targeted and untargeted assays in fasting plasma). GAD/IA-2 assessments from fasting serum samples were also undertaken. Data from the Recent Physical Activity Questionnaire (RPAQ) and sleep diaries were also collected in sub-cohorts.

### Statistical methods for descriptive data

Based on standard ADA criteria for glycaemic control [2], Cohort 1 was stratified into two categories: normal glucose regulation (NGR) and impaired glucose regulation (IGR). NGR was defined as having HbA1c, fasting glucose and 2-hr glucose values within the normal ranges for each measure. IGR was defined as having impaired values in at least one of HbA1c, fasting glucose or 2-hr glucose. Cohort 2 was stratified into treatment categories: lifestyle advice only (LS) or metformin plus lifestyle advice (LS+MET). Descriptive statistics are presented as mean±SD. Pairwise Pearson correlations were carried out between all key variables described here. For these analyses, continuous variables were first inverse normal transformed and then adjusted for age, sex, and study centre by two-step residual regression. We described the same for anthropometric and glycaemic control variables for the first main follow-up visit, as well as the difference between the baseline and follow-up visit (Follow-upΔ= follow-up value – baseline value). We also calculated pairwise Pearson correlations for the follow-upΔ values; for these analyses, continuous variables were first inverse normal transformed and then adjusted for age, sex, study centre and days since baseline visit by two-step residual regression. All statistics were computed using *R* software version 3.2.3 [21]. The IMI DIRECT data release version used for the analyses in this article was *direct*_*03*-*11*-*2017*.

### Glycaemic trait modelling

Glycemic traits were derived from frequently-sampled (0, 15, 30, 45, 60, 90, 120 min) 75g oral glucose tolerance tests (fsOGTT) or mixed-meal tolerance tests (MMTT; 0, 30, 60, 90, 120 min) for Cohort 1 and Cohort 2, respectively. Analyses used a mathematical model that describes the relationship between insulin secretion and glucose concentration [22, 23]. The model expresses insulin secretion as the sum of two components: the first component represents the dependence of insulin secretion on absolute glucose concentration at any time during the fsOGTT/MMTT, through a dose-response function. Characteristic parameters of the dose-response relationship are the mean slope over the observed glucose range, denoted as *glucose sensitivity.* The dose-response relationship is modulated by a potentiation factor, which accounts for the fact that during acute stimulation, insulin secretion is higher on the descending phase of hyperglycaemia than at the same glucose concentration on the ascending phase. In participants with normal glucose regulation and insulin secretion, the potentiation factor typically increases from baseline to the end of a 2-hr OGTT [24]. To quantify this excursion, the ratio between the 2-hr and the baseline values were calculated. This ratio is denoted as *potentiation ratio* and reflects late insulin release. The second insulin secretion component represents the dependence of insulin secretion on the rate of change of glucose concentration. This component is termed *derivative component*, and is determined by a single parameter, denoted as *rate sensitivity*. Rate sensitivity reflects early insulin release [24].

The model parameters were estimated from glucose and C-peptide concentrations by regularized least-squares, as previously described [22]. Regularization involves the choice of smoothing factors, which were selected to obtain glucose and C-peptide model residuals with standard deviations close to expected measurement error (~1% for glucose and ~4% for C-peptide). Insulin secretion rates were calculated from the model every 5 minutes. The integral of insulin secretion during the fsOGTT was denoted as total insulin output.

## Results

### Cohort 1 – Prediabetes

Of 2235 enrolled participants in Cohort 1, 2127 passed all inclusion, exclusion, and quality control criteria. Of these, 1419 (67%) had IGR according to at least one ADA category for HbA1c, fasting glucose or 2-hr glucose [2], and were thus within the target ‘prediabetic’ range. A total of 693 participants (33% of Cohort 1) had NGR for all three glycaemic measures. Participants with prevalent T2D (n=105) or who withdrew from the study (n=3) were excluded from the study.

The number of participants enrolled into Cohort 1 varied between centres, with the Finnish sub-cohort being the largest with 58% (n=1240) of the total Cohort 1 baseline sample. The other centres in the Netherlands, Denmark and Sweden enrolled 22% (n=473), 13% (n=275), and 7% (n=139) of the total cohort, respectively.

The ratio of men to women varied in each sub-cohort, with all participants at the Finnish centre being male, and 29%, 45% and 43% being male in the sub-cohorts from Sweden, the Netherlands and Denmark, respectively.

Detailed participant characteristics for baseline variables for Cohort 1, are shown in Table 1 (and stratified by glycaemic category in Supplementary Table 1). Figure 2 shows the pairwise correlations between a selection of key phenotypic variables at baseline. Participant characteristics at the 18-month follow-up and the change (delta) between baseline and follow-up for Cohort 1 are shown in Table 2. The pairwise correlations between the baseline-follow-up difference for anthropometric and glucose control variables are shown in Figure 3.

**Figure 2.**
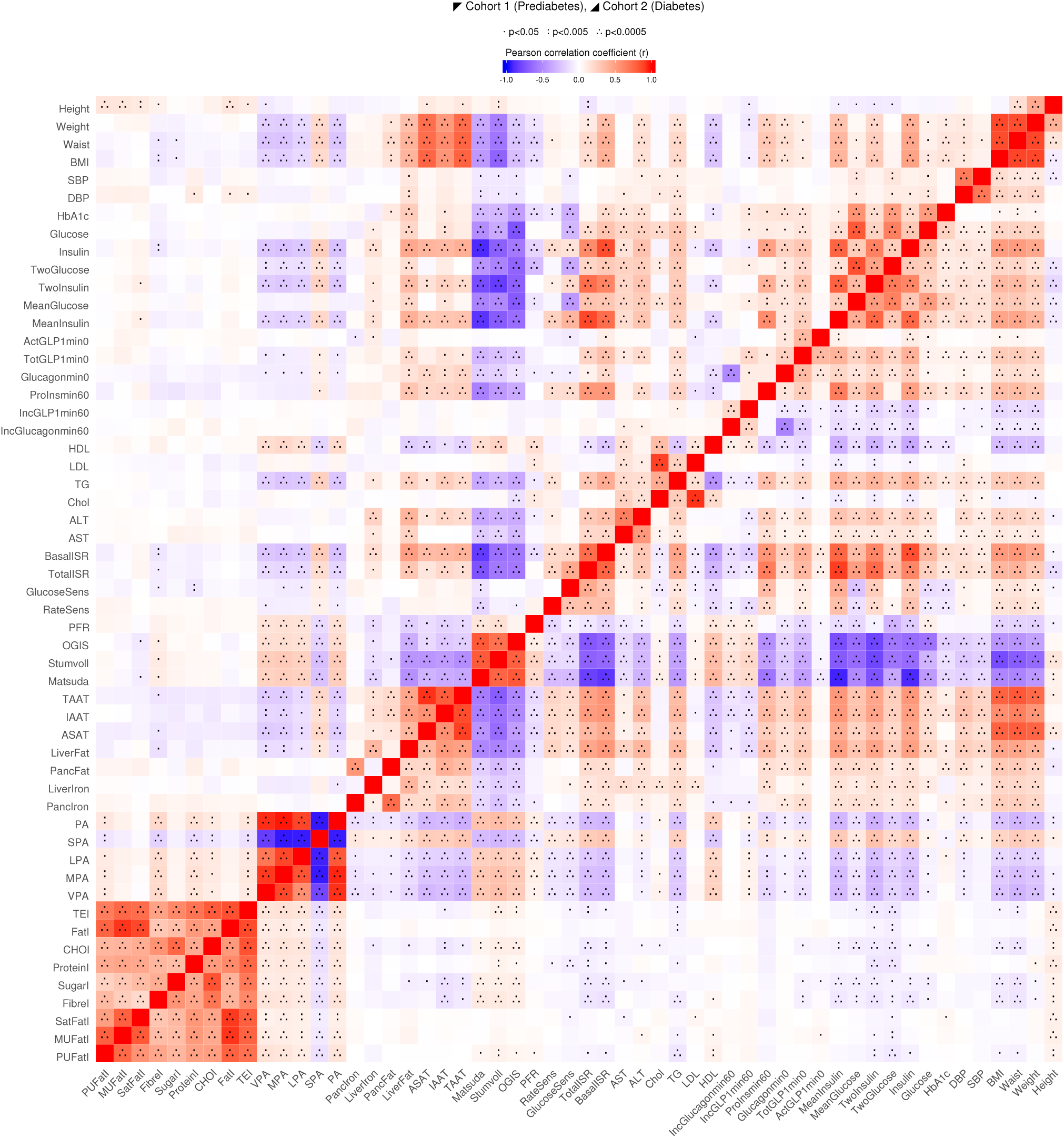
Pairwise correlation matrix. Fill colour indicates Pearson correlation coefficient (r), where positive is denoted by red fill, inverse by blue fill, magnitude by intensity. Cohort 1 and 2 are separate, above and below diagonal, respectively. All continuous variables were normally transformed and adjusted for age, sex, and study center. SBP: Systolic blood pressure, DBP: Diastolic blood pressure, Glucose: Fasting glucose, Insulin: Fasting insulin, TwoGlucose: 2-hr glucose, TwoInsulin: 2-hr insulin, MeanGlucose: Mean 2-hr glucose, MeanInsulin: Mean 2-hr insulin, ActGLP1min0: Fasting intact GLP-1 concentration, TotGLP1min0: Fasting total GLP-1 concentration, Glucagonmin0: Fasting glucagon, ProInsmin60: 1h intact proinsulin, IncGLP1min60: 1h GLP-1 increment, IncGlucagonmin60: 1h glucagon increment, HDL: Fasting HDL cholesterol, LDL: Fasting LDL cholesterol, TG: Fasting triglycerides, Chol: Total cholesterol, ALT: Alanine aminotransferase, AST: Aspartate transaminase, BasalISR: Fasting insulin secretion, TotalISR: Integral of total insulin secretion, GlucoseSens: Glucose sensitivity, RateSens: Rate sensitivity, PFR: Potentiation factor ratio, OGIS: Insulin sensitivity 2-h OGIS, Stumvoll: Insulin sensitivity Stumvoll, Matsuda: Insulin sensitivity Matsuda, IAAT: Intrabdominal Adipose Tissue, ASAT: Abdominal Subcutaneous Adipose Tissue, TAAT: Total Abdominal Adipose Tissue, LiverFat: Liver Fat, PancFat: Pancreatic Fat, LiverIron: Liver Iron content, PancIron: Pancreatic Iron content, PA: Average physical activity intensity - hpfVM, SPA: Sedentary, LPA: Light, MPA: Moderate, VPA: Vigorous, TEI: Total energy intake, FatI: Fat intake, CHOI: Carbohydrate intake, ProteinI: Protein intake, SugarI: Sugar intake, FibreI: Fibre intake, SatFatI: Saturated fat intake, MUFatI: Monunsaturated fat intake, PUFatI: Polyunsaturated fat intake.

**Figure 3.**
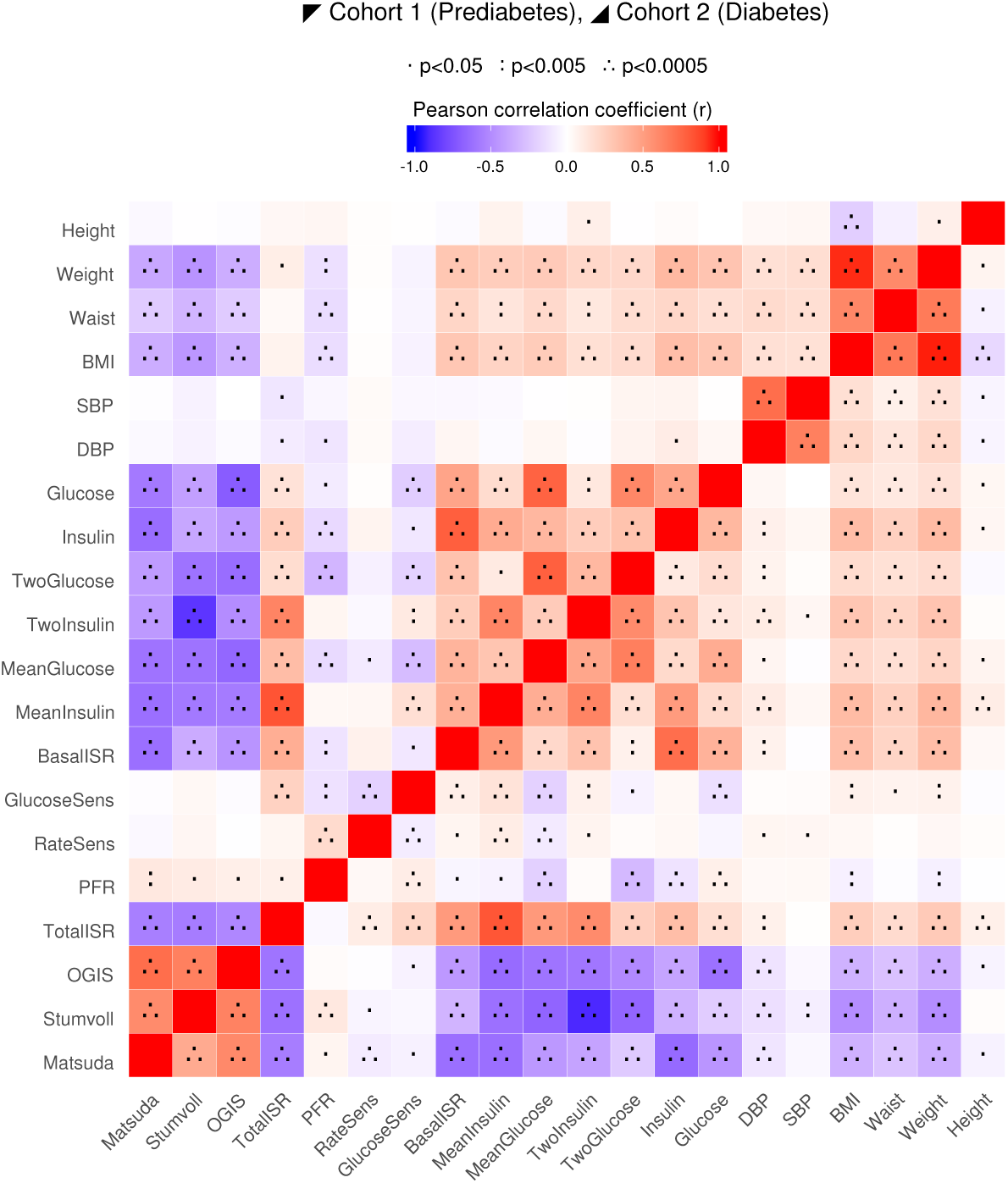
Pairwise correlation matrix of Follow-up **Δ**. (difference between characteristic value between follow-up assessment and baseline visits). Fill colour indicates Pearson correlation coefficient (r), where positive is denoted by red fill, inverse by blue fill, magnitude by intensity. Cohort 1 and 2 are separate, above and below diagonal, respectively. All continuous variables were normally transformed and adjusted for age, sex, and study center. SBP: Systolic blood pressure, DBP: Diastolic blood pressure, Glucose: Fasting glucose, Insulin: Fasting insulin, TwoGlucose: 2-hr glucose, TwoInsulin: 2-hr insulin, MeanGlucose: Mean 2-hr glucose, MeanInsulin: Mean 2-hr insulin, BasalISR: Fasting insulin secretion, TotalISR: Integral of total insulin secretion, GlucoseSens: Glucose sensitivity, RateSens: Rate sensitivity, PFR: Potentiation factor ratio, OGIS: Insulin sensitivity 2-h OGIS, Stumvoll: Insulin sensitivity Stumvoll, Matsuda: Insulin sensitivity Matsuda.

**Table 1.** Baseline clinical and phenotypic characteristics of Cohorts 1 and 2. Descriptive statistics shown are N, % or mean (standard deviation) as indicated. Values are untransformed and unadjusted.

|  | Cohort 1 (Prediabetes) |  | Cohort 2 (Diabetes) |  |
| --- | --- | --- | --- | --- |
|  | Mean (SD) | N | Mean (SD) | N |
| Male (%) | 76% | 2127 | 58% | 789 |
| Months since screening visit | 6.4(4.8) | 2127 | 0.9(0.9) | 787 |
| Age (yrs) | 62(6.2) | 2127 | 62(8.1) | 787 |
| Height (cm) | 174(8) | 2127 | 171(9.8) | 787 |
| Weight (kg) | 85(13) | 2127 | 89(17) | 787 |
| Waist circumference (cm) | 99(11) | 2127 | 103(13) | 781 |
| BMI (kg·m<sup>-2</sup>) | 27.9(4.0) | 2127 | 30.5(5.0) | 787 |
| Systolic blood pressure (mmHg) | 131(15) | 2107 | 131(16) | 664 |
| Diastolic blood pressure (mmHg) | 81(9) | 2107 | 75(9.5) | 664 |
| HbA1c (mmol·mol<sup>-1</sup>) | 37(2.9) | 2113 | 46(5.8) | 784 |
| Fasting glucose (mmol·L<sup>-1</sup>) | 5.7(0.6) | 2126 | 7.2(1.4) | 787 |
| Fasting insulin (pmol·L<sup>-1</sup>) | 10.9(7.6) | 2124 | 106.6(70.9) | 787 |
| Fasting HDL cholesterol (mmol·L<sup>-1</sup>) | 1.3(0.4) | 2123 | 1.2(0.4) | 789 |
| Fasting LDL cholesterol (mmol·L<sup>-1</sup>) | 3.2(0.9) | 2123 | 2.3(1) | 781 |
| Fasting triglycerides (mmol·L<sup>-1</sup>) | 1.4(0.6) | 2123 | 1.5(0.9) | 789 |
| Alanine aminotransferase (IU·L<sup>-1</sup>) | 18(12) | 2120 | 26(14) | 789 |
| Aspartate transaminase (IU·L<sup>-1</sup>) | 27(10) | 2052 | 26(12) | 789 |
| Total cholesterol (mmol·L<sup>-1</sup>) | 5.1(1) | 2123 | 4.2(1.2) | 789 |
| Fasting intact GLP-1 concentration (pg·mL<sup>-1</sup>) | 0.41(0.59) | 2121 | 0.67(1.05) | 782 |
| Fasting total GLP-1 concentration (pg·mL<sup>-1</sup>) | 6.5(4.4) | 2120 | 9.4(9) | 780 |
| Fasting glucagon (pg·mL<sup>-1</sup>) | 98(41) | 2116 | 111(51) | 758 |
| 1hr intact proinsulin (pg·mL<sup>-1</sup>) | 19(11.7) | 578 | 21(13.6) | 382 |
| 1hr GLP-1 increment (pg·mL<sup>-1</sup>) | 9.3(12.1) | 2103 | 9.8(12.5) | 774 |
| 1hr glucagon increment (pg·mL<sup>-1</sup>) | -10.7(38) | 2097 | -3.9(51) | 746 |
| Mean 2hr glucose (mmol·L<sup>-1</sup>) | 7.7(1.5) | 2126 | 9.3(2) | 779 |
| Mean 2hr insulin (pmol·L<sup>-1</sup>) | 383(260) | 2126 | 457(275) | 779 |
| 2hr glucose (mmol·L<sup>-1</sup>) | 5.9(1.6) | 2127 | 8.6(2.8) | 786 |
| 2hr insulin (pmol·L<sup>-1</sup>) | 48(48) | 2102 | 445(348) | 786 |
| Fasting insulin secretion (pmol·min<sup>-1</sup>·m<sup>-2</sup>) | 106(40) | 2126 | 137(48) | 779 |
| Integral of total insulin secretion (nmol·m<sup>-2</sup>) | 52(18) | 2126 | 44(14) | 779 |
| Glucose sensitivity (pmol·min<sup>-1</sup>·m<sup>-2</sup>·mmol<sup>-1</sup>·L<sup>-1</sup>) | 113(55) | 2126 | 83(55) | 779 |
| Rate sensitivity (pmol·m<sup>-2</sup>·mmol<sup>-1</sup>·L<sup>-1</sup>) | 921(699) | 2126 | 1124(1082) | 779 |
| Potential factor ratio (dimensionless) | 1.7(0.6) | 2126 | 1.4(0.6) | 777 |
| Insulin sensitivity 2hr OGIS (ml·min<sup>-1</sup>·m<sup>-2</sup>) | 381(59) | 2118 | 298(69) | 775 |
| Insulin sensitivity Stumvoll (ml·min<sup>-1</sup>·kg<sup>-1</sup>) | 7.8(2.4) | 2099 | 5.5(2.7) | 775 |
| Insulin sensitivity Matsuda (arbitrary) | 5(3.1) | 2126 | 2.9(2.2) | 779 |
| Intrabdominal Adipose Tissue (L) | 5.5(2.4) | 956 | 5.7(2.2) | 374 |
| Abdominal Subcutaneous Adipose Tissue (L) | 6.1(2.6) | 953 | 8.1(3.8) | 374 |
| Total Abdominal Adipose Tissue (L) | 12(3.9) | 953 | 14(4.8) | 374 |
| Liver Fat (%) | 5(4.7) | 959 | 8.7(7.1) | 498 |
| Pancreatic Fat (%) | 13(8.9) | 929 | 11(7.3) | 446 |
| Liver Iron content (mg·g<sup>-1</sup>) | 1.3(0.26) | 958 | 1.4(0.31) | 498 |
| Pancreatic Iron content (mg·g<sup>-1</sup>) | 1.3(0.43) | 927 | 1.2(0.33) | 447 |
| Average physical activity intensity - hpFVM (mGs) | 37(10.2) | 1714 | 34(9.9) | 722 |
| Sedentary (% of time) | 82(4.2) | 1714 | 83(4.3) | 722 |
| Light (% of time) | 10.9(2.3) | 1714 | 10.4(2.3) | 722 |
| Moderate (% of time) | 5.3(1.5) | 1714 | 4.9(1.6) | 722 |
| Vigorous (% of time) | 1.5(0.7) | 1714 | 1.3(0.7) | 722 |
| Protein (g·day<sup>-1</sup>) | 1963(751) | 2064 | 1840(602) | 707 |
| Carbohydrate (g·day<sup>-1</sup>) | 223(96) | 2064 | 213(78) | 707 |
| Fat intake (g·day<sup>-1</sup>) | 79(39) | 2064 | 72(33) | 707 |
| Total Energy (kCal·day<sup>-1</sup>) | 99(44) | 2064 | 87(31) | 707 |
| Sugar (g·day<sup>-1</sup>) | 96(53) | 2064 | 85(43) | 707 |
| Fibre (g·day<sup>-1</sup>) | 20(9.8) | 2064 | 19(8.4) | 707 |
| Saturated fat (g·day<sup>-1</sup>) | 29(16) | 2064 | 26(14) | 707 |
| Monounsaturated fat (g·day<sup>-1</sup>) | 27(17) | 2064 | 24(13) | 707 |
| Polyunsaturated fat (g·day<sup>-1</sup>) | 13(8.2) | 2064 | 12(8) | 707 |

**Table 2.** Follow-up and Follow-up **Δ** in clinical and phenotypic characteristics of Cohorts 1 and 2. IGR: Impaired glucose regulation, NGR: Normal glucose regulation, LS: Lifestyle Treatment, LS+MET: Lifestyle and metformin treatment, Descriptive statistics shown are N, % or mean (standard deviation) as indicated. Values are untransformed and unadjusted. Follow-up **Δ** is mean (sd) of difference between characteristic value between follow-up assessment and baseline visits.

|  | Cohort 1 (Prediabetes) |  |  | Cohort 2 Diabetes) |  |  |
| --- | --- | --- | --- | --- | --- | --- |
| | Follow-up<br>Mean (SD) | Follow-up $\Delta$<br>Mean (SD) | N | Follow-up<br>Mean (SD) | Follow-up $\Delta$<br>Mean (SD) | N |
| Male (%) | 77% |  | 1989 | 60% |  | 668 |
| Months since baseline visit | 18.6(1.4) |  | 1989 | 18.2(0.6) |  | 668 |
| Age (yrs) | 64(6.1) |  | 1989 | 64(7.7) |  | 668 |
| Height (cm) | 174(7.9) | -0.16(0.76) | 1983 | 171(9.8) | -0.1(1.22) | 662 |
| Weight (kg) | 85(14) | -0.18(3.6) | 1981 | 90(17) | 0.13(4.6) | 664 |
| Waist circumference (cm) | 99(11) | -0.3(4.7) | 1981 | 103(13) | 0.5(6.4) | 661 |
| BMI ($\text{kg}\cdot\text{m}^{-2}$) | 27.9(4.0) | -0.01(1.2) | 1980 | 30.5(5.0) | 0.09(1.6) | 661 |
| Systolic blood pressure (mmHg) | 131(16) | 0.1(13) | 1983 | 130(16) | -1.5(15) | 667 |
| Diastolic blood pressure (mmHg) | 80(8.8) | -0.7(7.1) | 1983 | 74(9.3) | -1.5(8.3) | 665 |
| Fasting glucose ($\text{mmol}\cdot\text{L}^{-1}$) | 5.8(0.6) | 0.1(0.5) | 1977 | 7.8(1.8) | 0.6(1.6) | 658 |
| Fasting insulin ($\text{pmol}\cdot\text{L}^{-1}$) | 11.8(8.6) | 1.1(6.3) | 1975 | 118.8(78.6) | 14.7(60.3) | 655 |
| 2hr glucose ($\text{mmol}\cdot\text{L}^{-1}$) | 6.1(1.7) | 0.2(1.6) | 1975 | 9.5(3.3) | 0.9(2.9) | 659 |
| 2hr insulin ($\text{pmol}\cdot\text{L}^{-1}$) | 55(54) | 7.3(38) | 1950 | 450(313) | 13.9(273.7) | 648 |
| Mean 2hr glucose ($\text{mmol}\cdot\text{L}^{-1}$) | 7.8(1.6) | 0.1(1.2) | 1974 | 10.1(2.6) | 0.8(2.1) | 649 |
| Mean 2hr insulin ($\text{pmol}\cdot\text{L}^{-1}$) | 425(280) | 42(171) | 1974 | 476(279) | 21(172) | 649 |
| Fasting insulin secretion ($\text{pmol}\cdot\text{min}^{-1}\cdot\text{m}^{-2}$) | 103(41) | -2.4(26) | 1974 | 145(52) | 8.9(35) | 649 |
| Glucose sensitivity ($\text{pmol}\cdot\text{min}^{-1}\cdot\text{m}^{-2}\cdot\text{mmol}^{-1}\cdot\text{L}^{-1}$) | 109(55) | -3.9(56) | 1974 | 81(58) | -3.2(53) | 649 |
| Rate sensitivity ($\text{pmol}\cdot\text{m}^{-2}\cdot\text{mmol}^{-1}\cdot\text{L}^{-1}$) | 833(673) | -83(698) | 1974 | 1176(1291) | 44(1404) | 649 |
| Potential factor ratio (dimensionless) | 1.8(0.7) | 0.1(0.8) | 1973 | 1.4(0.5) | 0(0.7) | 649 |
| Integral of total insulin secretion ($\text{nmol}\cdot\text{m}^{-2}$) | 50(19) | -1.7(13) | 1974 | 45(14) | 1.2(10.4) | 649 |
| Insulin sensitivity 2hr OGIS ($\text{ml}\cdot\text{min}^{-1}\cdot\text{m}^{-2}$) | 369(61) | -11(45) | 1963 | 278(54) | -20(61) | 640 |
| Insulin sensitivity Stumvoll ($\text{ml}\cdot\text{min}^{-1}\cdot\text{kg}^{-1}$) | 7.6(2.6) | -0.2(1.6) | 1941 | 5.3(2.7) | -0.3(1.9) | 639 |
| Insulin sensitivity Matsuda (arbitrary) | 4.7(3.1) | -0.3(2.2) | 1974 | 2.5(1.7) | -0.5(1.6) | 648 |

Briefly, at baseline, participants had a mean(SD) age of 62(6.2) years, BMI of 27.9(4.0) kg/m^2^, HbA1c of 37(2.9) mmol/mol, fasting glucose of 5.7(0.6) mmol/l, 2-hr glucose of 5.9(1.6) mmol/l, fasting insulin of 10.9(7.6) pmol/l, glucose sensitivity of 113(55) pmol min^-1^m^-2^mM^-1^, and insulin sensitivity (2-hr OGIS) of 381(59) ml min^-1^m^-2^.

### Cohort 2 - New-onset diabetes

Of 830 patients enrolled to attend the screening visit in Cohort 2, 789 passed all inclusion, exclusion, and quality control criteria. Of these, 271 were receiving LS+MET and 508 were LS. Participants who withdrew consent, were receiving any other oral hypoglycaemic agent, or having ever received insulin treatment were been excluded (n=41). Of those enrolled into Cohort 2, the UK (Dundee, Exeter, Newcastle), Dutch (Amsterdam), Swedish (Lund), and Danish (Copenhagen) study centres enrolled 21% (n=167), 18% (n=141), 21% (n=167), 20% (n=158), 12% (n=96) and 7% (n=52) of the total cohort, respectively; 52% to 63% of these sub-cohorts were male.

Detailed participant characteristics at baseline for key variables for Cohort 2 are shown in Table 1 (and stratified by treatment category in Supplementary Table 1). Figure 1 shows the pairwise correlations between key variables at baseline adjusted for age, sex and study centre in a correlation matrix. Participant characteristics for follow-up and the baseline-to-18m follow-up (change/delta) difference for Cohort 2 is shown in Table 2. Figure 3 shows, the pairwise Pearson correlations between the baseline-follow-up differences for anthropometric and glucose control variables.

Briefly, at baseline, participants had a mean(SD) age of 62(8.1) years, BMI of 30.5(5.0) kg/m^2^, HbA1c of 46.5(5.8) mmol/mol, fasting glucose of 7.2(1.4) mmol/l, 2-hr glucose of 8.6(2.8) mmol/l, fasting insulin of 107(71) pmol/l, glucose sensitivity of 83(55) pmol min^-1^m^-2^mM^-1^, and insulin sensitivity (2-hr OGIS) of 298(69) ml min^-1^m^-2^.

### Population substructure

As some study centres enrolled participants into both cohorts, we elected to characterise the population substructure across the cohorts by study centre (i.e., pooling both cohorts at a given centre where possible). As illustrated in Figure 4, genetic substructures closely map to the geographic location of the populations, indicating ethnic homogeneity within regions from which the cohorts were recruited, whereas there is far greater heterogeneity between centres, the latter driven mainly by the inclusion of Finnish participants.

**Figure 4.**
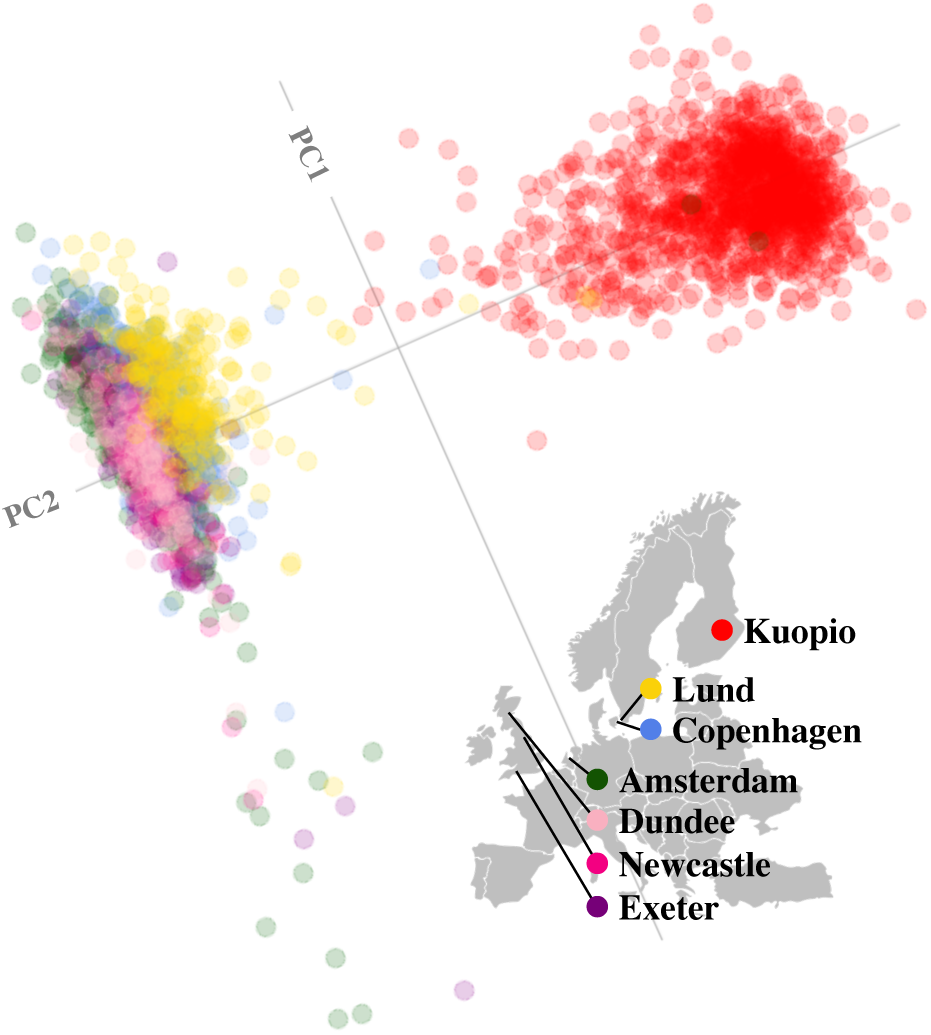
Population structure within WP2 baseline study samples. A statistical summary of genetic data from Cohorts 1 and 2 (combined) based on principal component axis one (PC1) and axis two (PC2). Points are coloured as per the recruitment centres. Red, Kuopio; Yellow, Lund; Blue, Copenhagen; Green, Amsterdam; Pink, Newcastle; Salmon, Dundee; Purple, Exeter.

## Discussion

Major advances in technologies and methods over the past decade make high-resolution quantification of disease phenotypes and processes possible in large sample collections. Applying modern assays to historical biosamples is particularly useful when studying processes that take decades to unfold. However, biosamples often degrade with long-term storage and many older studies did not deploy the advanced phenotyping methods available today. Recognizing these limitations, we designed and initiated two state-of-the-art prospective cohort studies as part of the IMI DIRECT consortium. Designed for biomarker discovery in glycaemic deterioration and diabetes progression, the IMI DIRECT cohorts include conventional and cutting-edge phenotyping techniques and technologies that are repeated on multiple occasions during a follow-up period of up to 48-months (currently ongoing). Here, we report the characteristics of the IMI DIRECT cohorts at baseline and 18-months follow-up for glycaemic deterioration, and consider these results in the context of the implemented protocols and the plans outlined in the design and rationale paper published previously [7].

The recruitment strategies for the two IMI DIRECT cohorts differed in that Cohort 1 focused on recruiting from an existing large sample-frame (N=24,682) derived from established prospective cohort studies, whereas Cohort 2 used clinical registries to identify eligible participants. The strategy for recruiting participants from existing prospective studies for Cohort 1 facilitated access to data that was used to predict risk of rapid glycaemic deterioration. However, despite the relatively large sampling-frame, it was necessary to enrol lower-risk participants, in order to achieve the target sample size; in doing so, we recognized that this would likely reduce the overall rate of glycaemic deterioration. In Cohort 2, we fell slightly short of the target sample size of 1000 participants (n=789 with complete and high-quality data eventually enrolled), which reflects difficulties faced in engaging general practices at some study sites.

The descriptive statistics, pairwise correlations and genetic substructures presented in this article are not intended to make aetiological inferences, but rather to compliment subsequent IMI DIRECT papers by providing details and context.

The two IMI DIRECT cohorts are distinct, albeit with many close parallels in methodology, allowing similar analyses to be carried out in both cohort datasets. Comparisons between Cohort 1 and Cohort 2 may help determine whether biomarkers for glycaemic deterioration are conditional on diabetes. However, some key differences in study design between cohorts (e.g. fsOGTT vs MMTT) may inhibit some comparisons. Differentiating true differences from error for almost all data shown here when only two time-points are available is challenging. Subsequent analyses, including assessments obtained from a third time-point (up to 48-months after baseline) will be available in the future, which will help determine the extent to which the delta values shown here reflect true change.

### Conclusion

The study described here is being used to unravel the heterogeneous pathophysiology of prediabetes and diabetes and to discover biomarkers that might prove useful for patient stratification and therapeutic optimization. As more prospective data are accrued, the IMI DIRECT cohorts will grow in value. In the long-term, the IMI DIRECT consortium intends to make these data available to other researchers through a managed-access repository.

## Acknowledgements

We thank all the participants and study centre staff in IMI DIRECT for their contribution to the study.

